# Genetic Characterization of the Cell Types of in Developing Feathers, and the Evolution of Feather Complexity

**DOI:** 10.1101/2024.11.21.624532

**Authors:** Cody Limber, Günter P. Wagner, Richard O. Prum

## Abstract

Feathers are the most complex and diverse epidermal appendages found in vertebrates. Their unique hierarchical organization and development is based on a diversity of cell types and morphologies. Despite being well characterized morphologically and extensive molecular developmental research focusing on candidate genes, little is known about the gene regulatory identities of these presumptive feather cell types. Here, we use single cell and single nuclear RNA sequencing with *in situ* hybridization to identify and characterize cells types in embryonic chicken feathers. We show that the distinct cell morphologies correspond to feather cell types with distinct gene expression profiles. We also describe a previously unidentified cell type, the basal barb ridge epithelium, which appears to play a role in signaling necessary for barb ridge differentiation and pulp cap production. We also analyze RNA velocity trajectories of developing feather cells, and find distinct developmental trajectories for epidermal cells that constitute the mature feather and those that function only in feather development. Finally, we produce an evolutionary tree of feather cell types based on transcription factor expression in order to test prior developmental hypotheses about feather evolution. Our tree is consistent with the developmental model of feather evolution, and sheds light on the influence of ancestral epidermal stratification on feather cell evolution. This transcriptomic approach to study feather cell types helps lay the ground work for understanding the developmental evolutionary complexity and diversity of feathers.

## Introduction

Advances in single cell sequencing technologies have created new intellectual connections between molecular genetics, morphological development, and evolutionary biology.^1^ While previous studies have documented the vital role of development in the evolution of anatomical innovations, molecular characterization of the transcriptomic states of diverse cell types within evolutionary novelties provides new opportunities to investigate the development and evolution of hierarchically complex anatomical structures.^2–5^

An exemplar evolutionary novelty is the avian feather. Feathers are branched epidermal appendages with extraordinary morphological diversity within and among feather tracts, plumages, and bird species. Feathers are critical to many aspects of avian biology, including thermal homeostasis, water repellency, flight, coloration, camouflage, display, and even sound production.^6–15^ In addition to the diversity among bird species, feathers also vary across the body of individual birds, and over the course of a bird’s development, from a fluffy chick’s natal down to an adult’s pennaceous flight feathers. Characterizing feather cell types from transcriptomic data can help us better understand the genetic mechanisms involved in the development of innovations unique to feathers, and contribute to our understanding of the evolution of feather complexity and diversity.^6,16^

Feathers present a particularly interesting system for the investigation of the development and evolution of cell types. There is a rich classical literature describing the hierarchical, modular development of feather morphology.^6,17^ (For a more comprehensive introduction to feather structure and development, see the Supplementary Information). Since the publication of a developmental model of feather evolution in 1999,^3^ feathers have become an important system for the investigating molecular development and evolution.^16,18–22^

Lucas and Stettenheim’s (1972; Figure 241) classic summary of feather development provides heuristic predictions of the possible differentiated cell types of the feather and feather germ.^6^ To identify and genetically characterize the cell types of the developing embryonic feather germ, we used transcriptomics of single cells and nuclei from plucked embryonic feathers and skin of leghorn chicken embryos. We identified clusters from our single cell data, and used in situ hybridization to verify these genetic clusters morphologically. We also used RNA velocity to hypothesize ontogenetic relationships among cell types, and produced a hypothesized tree of feather cell type evolution based on transcription factor expression. Our results suggest a model of feather evolution based on a stepwise addition of specialized cell types responsible for the key morphological and developmental innovations, leading to the complex morphology and diversity of extant feathers.

## Results

We sequenced transcriptomes from four samples – two single cell, and two single nuclear – of tissues collected from the spinal tract of chicken embryos over Hamburger-Hamilton stages 38-40 (Table 1). Three of the four samples included multiple, plucked feather germs, and one sample included multiple feathers with associated skin. We sequenced 5,715 cells from our skin and feather sample, and 14,684 cells from our three plucked feather samples. We identified 21 cell clusters in the aggregated data set from our plucked feather samples (Figure 1A), and 18 in our feathers-and-skin sample (Figure S1). We identified 13 epidermal cell clusters in a more focused analysis of epidermal cells isolated from the 3 plucked feather samples (Figure 2A). We mapped cell transcriptomic clusters to morphological cell types by *in situ* hybridization staining of cluster-identifying marker genes (Figure 1B-Q, 2B-DD). We also mapped cell development though an RNA velocity analysis (Figure 3), and proposed an evolutionary tree of feather cell types based on transcription factor expression (Figure 4).

**Figure 1:**
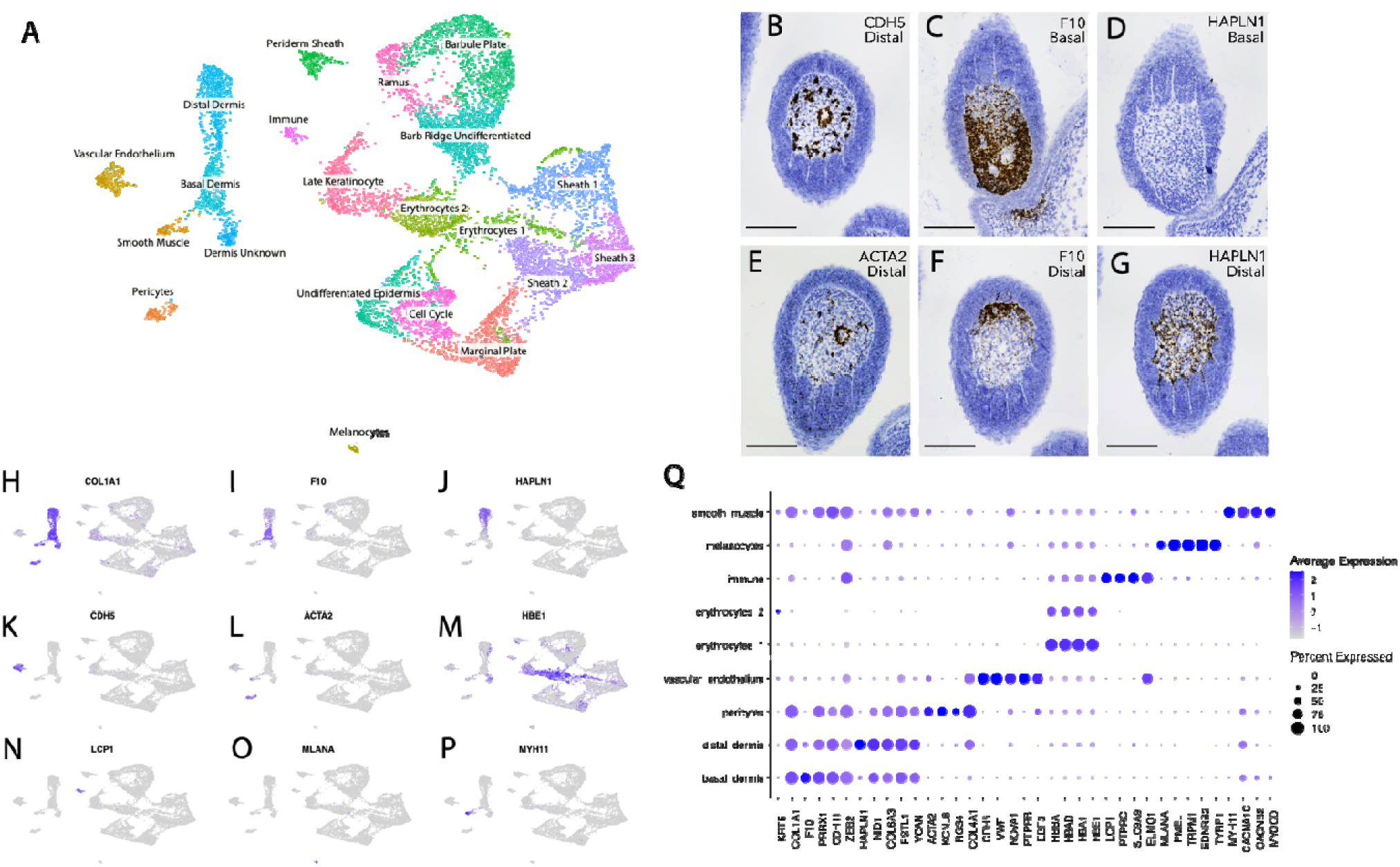
Single cell and nuclear sequencing identification of feather pulp cell types. (A) Uniform Manifold Approximation and Projection (UMAP) plot of all cells from plucked feather samples (Table 1). 21 cell types are color coded, and labeled with proposed cell type identities. (B-G) In situ hybridization stained histological sections for feather pulp cell type marker genes. For F10 and HAPL1, C and D are from the base of the feather germ, and F and G are from nearer the distal tip. Scale bars 0.1mm. (H-P) Feature plots highlighting expression patterns for specific marker genes used in feather pulp cell type identification. (Q) Dotplot of selected marker gene expression levels of embryonic feather pulp cell types.

**Figure 2:**
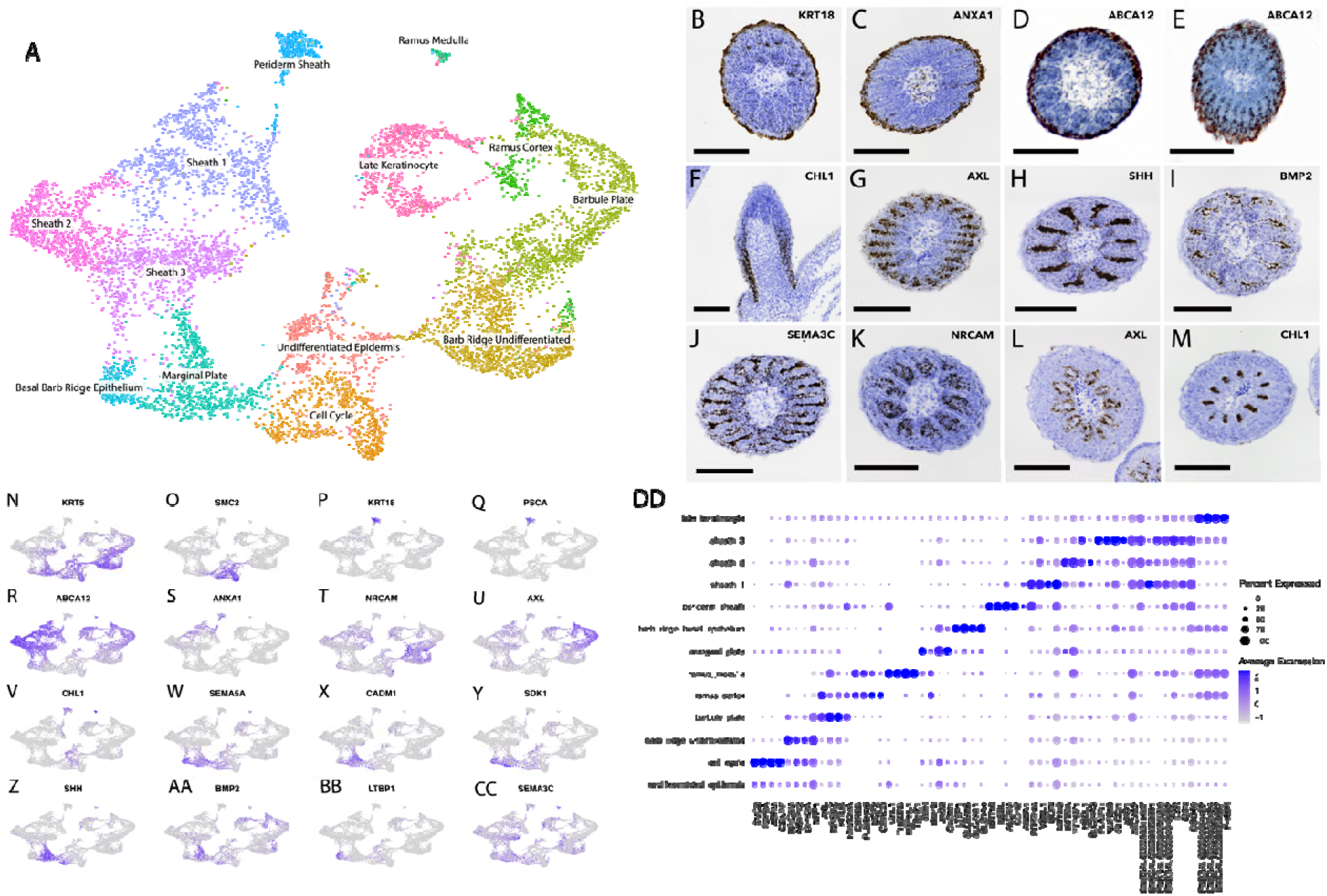
Analysis of epidermal cells reveals a diversity of feather epidermal cell types. (A) UMAP plot of epidermal cells colored and labeled by cell type identity. The ramus cell types and basal epithelium cell types were sub-clustered to identify rare cell types contained within those clusters. (B-M) In situ hybridization stained histological sections of genes used in epidermal cell type identification. See Supplemental Figure 4A and 4B for a close up of B and C. Scale bars 0.1mm (N-CC) Feature plots highlighting expression levels for specific marker genes used in epidermal cell type cluster identification. (DD) Dot plot of selected gene expression levels for marker genes for epidermal feather cell types

**Figure 3:**
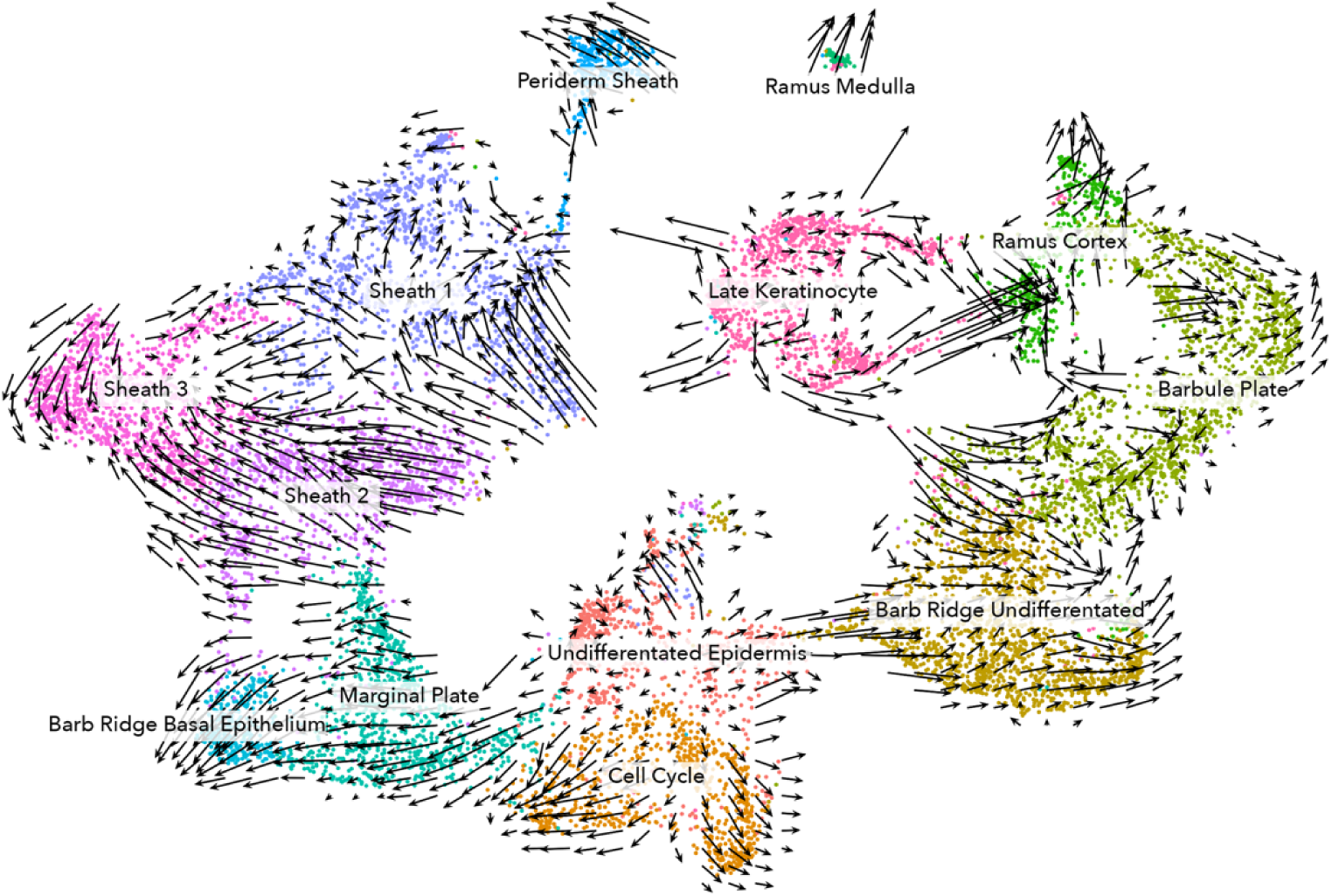
RNA velocity analysis mapped onto the UMAP plot of epidermal cell types. Arrows indicate the extrapolated future state of the cells in this region of the plot.

**Figure 4:**
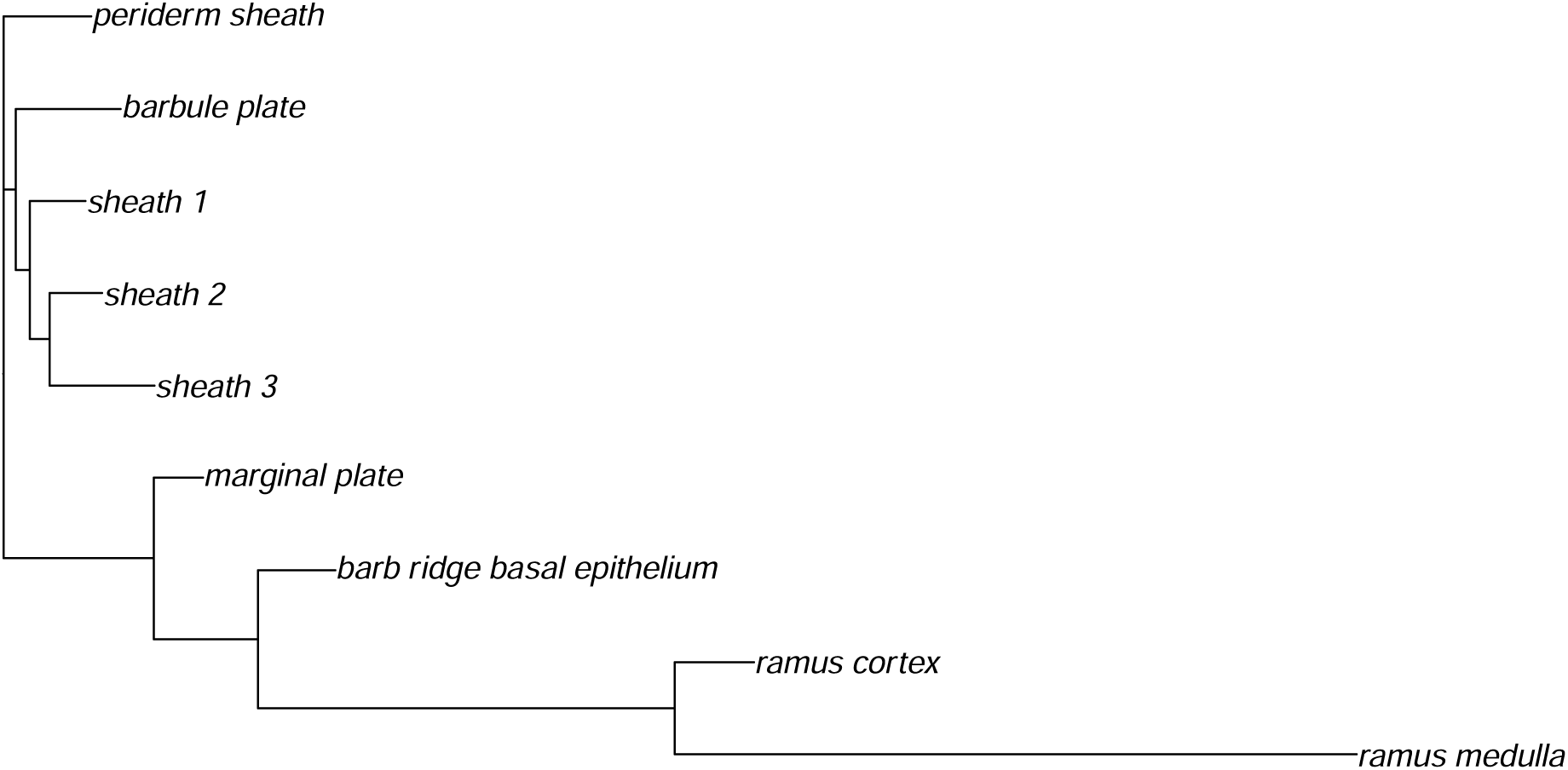
A neighbor-joining tree of feather germ cell types constructed from averaged transcription factor expression for each cell type cluster, rooted with periderm sheath.

**Table 1:** The developmental stage, age, sequencing method, and feather developmental stage for the four chick spinal skin samples sequenced. Developmental stages described indicate the most advanced morphologies developed in that sample (See SI2).

|  | H&H Stage | Sample type | Days Post Fertilization | Barbule Initiation Stage | Ramus Development Stage |
| --- | --- | --- | --- | --- | --- |
| Feather + Skin | 38 | Single cell | 12 | X |  |
| Plucked Feather | 38 | Single cell | 12 | X |  |
| Plucked Feather | 39 | Single nucleus | 13 |  | X |
| Plucked Feather | 40 | Single nucleus | 14 |  | X |

**Table 2:**
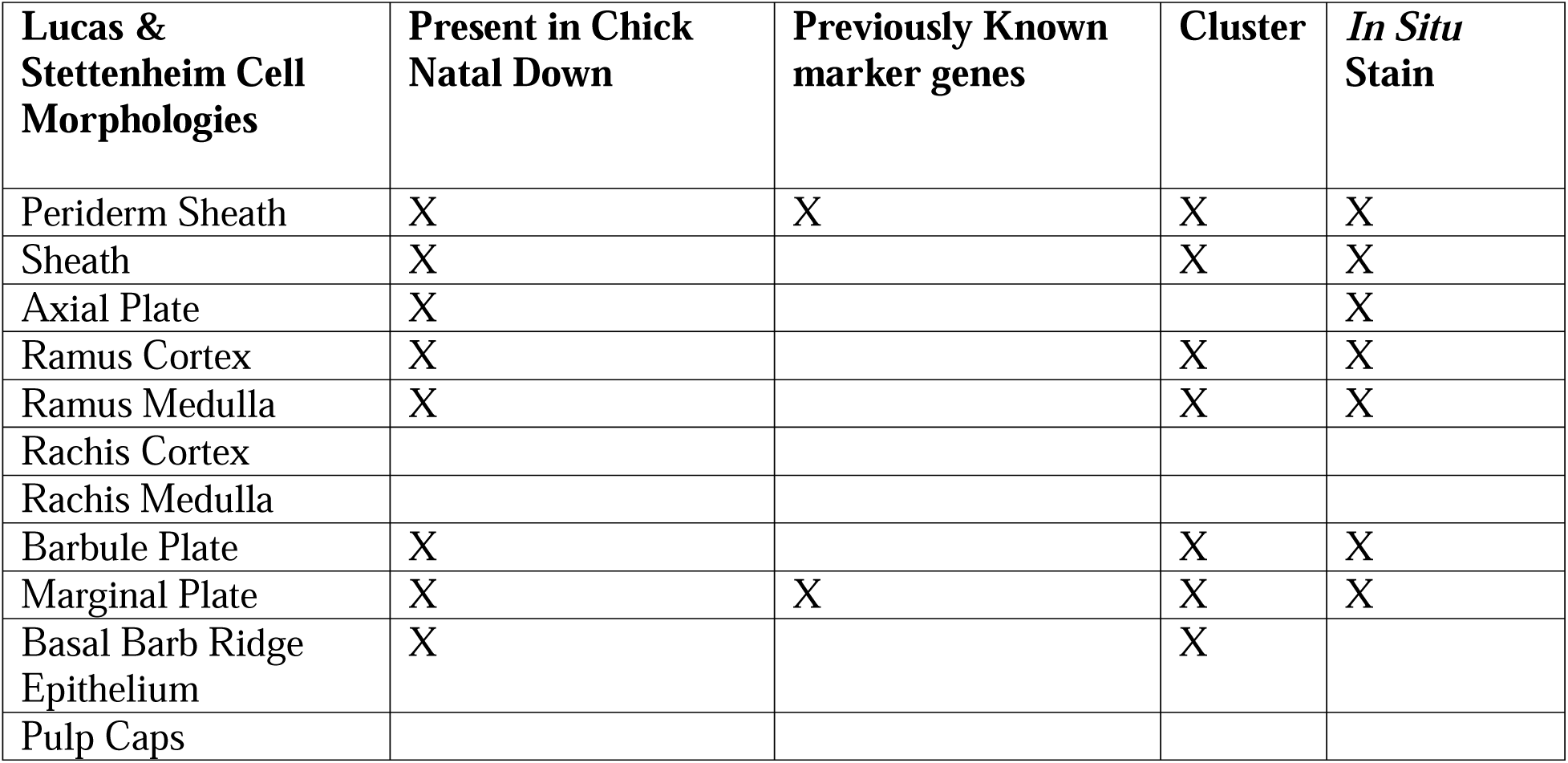
A summary of the epidermal cell morphologies described by Lucas and Stettenheim (Figure 241),^6^ their presence or absence in chicken natal down feathers, and our method of identification.

| **Lucas & Stettenheim Cell Morphologies** | **Present in Chick Natal Down** | **Previously Known marker genes** | **Cluster** | ***In Situ* Stain** |
| --- | --- | --- | --- | --- |
| Periderm Sheath | X | X | X | X |
| Sheath | X |  | X | X |
| Axial Plate | X |  |  | X |
| Ramus Cortex | X |  | X | X |
| Ramus Medulla | X |  | X | X |
| Rachis Cortex |  |  |  |  |
| Rachis Medulla |  |  |  |  |
| Barbule Plate | X |  | X | X |
| Marginal Plate | X | X | X | X |
| Basal Barb Ridge Epithelium | X |  | X |  |
| Pulp Caps |  |  |  |  |

### Pulp Cell Types

The feather pulp forms the center of the tubular feather germ, and it contains a complex cluster of cells (Figure 1) characterized by the expression collagen type I alpha 1 (*COL1A1*) (Figure 2), which is a major component of the dermal extracellular matrix.^23^ This cluster consists of dermal fibroblast, and contains three sub-clusters. The first sub-cluster (Figure 1A) differentially expresses *F10*, *PRRX1*, *CDH11*, and *ZEB2* (Figure 1I, Q). *F10* (coagulation factor X) is a gene that functions in blood coagulation,^24^ and its expression is localized to fibroblasts at the base of the feather follicle at all stages of development (Figure 1C, F). The second sub-cluster (Figure 1A) differentially expresses *HAPLN1*, *NID1*, *COL6A3*, and *FSTL1* (Figure 1J, Q). *HAPLN1* (hyaluronan-and-proteoglycan-link-protein 1), which is a component of the extracellular matrix and other tissues,^25,26^ localized to the dermal fibroblasts in the ramogenic zone distal to the *F10* expressing fibroblasts (Figure 1D, G). We suggest that these two dermal sub-clusters consist of temporal developmental states within a single dermal pulp fibroblast cell type (See Lucas and Stettenheim 1972; p. 365).^6^ The third sub cluster (Figure 1A) did not strongly differentially express any genes that were not also expressed by the other two clusters and thus, may be a transitory cell population.

In addition to dermal fibroblasts, the pulp includes a variety of other cell types that lack both keratin and *COL1A1* expression. One cluster of cells (Figure 1A) expresses *CDH5*, *VWF*, *NOVA1*, *PTPRB*, and *EBF3* (Figure 1K, Q). *VWF* is a known endothelial marker gene^27^ and *CDH5* (cadherin 5), which plays a role in development and maintenance of endothelial adherens junctions,^28^ is localized to the vascular endothelial cells around the pulp vasculature and axial arteries (Figure 1B).

A separate and distinct pulp cell cluster (Figure 1A) expresses smooth muscle α-actin *ACTA2* (Figure 1L), which is involved in maintaining cellular structure and integrity but is known for its role in cell contraction in myofibroblasts, smooth muscle, and pericytes.^29,30^ This cluster also expresses *KCNJ8* and *RGS4* (Figure 1Q), which localize to pericytes in human lung and heart.^31^ Staining for *ACTA2* (Figure 1E) also localizes this cell type to the perivascular tissue around the pulp vasculature and axial artery, but this distinct expression profile identifies these cells as pericytes.^29,32^

A small cluster of cells (Figure 1A) expresses a variety of smooth muscle marker genes including *MYH11*, *CANA1C*, and *MYOCD* (Figure 1P, Q). These cells are apparently part of the erector pili muscles that connect to the base of the feather and were pulled from the skin with the feather as it was plucked. We find a larger population of these smooth muscle cells in the sample of feather and skin (Figure S1).

We identified a pair of cell clusters (Figure 1A) that express high levels of hemoglobin genes, such as *HBBA*, *HBAD*, *HBA1*, and *HBE1* (Figures 1C and 1D). We identified these cells as erythrocytes. Another cluster of cells (Figure 1A) expressing lymphocyte cytosolic protein 1 (*LCP1*) and the pan-leukocyte marker *PTPRC*, also known as CD45 (Figures 1C and 1D), and were thus identified as immune cells.^33^

A small, highly distinct cluster of cells (Figure 1A) express a suite of genes associated with melanogenesis, including melan-A (*MLANA*) and premelanosome protein (*PMEL*) (Figure 1O, Q). Although leghorn chicks and chickens lack melanized feathers, we identified these cells as melanocytes, which are known to migrate into the dermal pulp of developing feather germs.^34–36^

### Epidermal Cell Types

Feather epidermal cells are diagnosable from pulp cells by the abundant expression α- and β-keratins. Early in feather cell development, keratin expression is dominated by α-keratins, including α-keratin 5 (*KRT5*), which we used to distinguish epidermal cells from pulp cells. Expression of (*KRT5*) diminishes as feather cells mature, and is replaced by expression of β-keratins.^37^

As we would predict from histological studies,^6^ feather epidermal cells contain the greatest heterogeneity of cells types in the feather germ. We identified 13 clusters of epidermal cells (Figure 2A). Furthermore, we find both fully differentiated cell types identified by localization of marker genes to mature cell morphologies, and intermediate developmental states identified by a transient gene expression profile, that are superseded by later developmental states, or cell types.

A cluster of epidermal cells (Figure 2A) expressing *CHL1*, (Figure 2V, DD) localized to the undifferentiated epidermis at the base of the feather germ (Figure 2F). This appears to be a transitory epidermal cell state between the undifferentiated feather epidermis and the fully differentiated cell types of the barb ridge that is not maintained once a distinct sheath or barb ridges develop.

Another cluster of epidermal of cells (Figure 2A) was diagnosed by expression of *SMC2* (Figure 2O, DD) which is involved in chromosome condensation during mitosis.^38^ Also expressed by this cluster are *ASPM,* which is involved in mitotic spindle construction,^39^ and *CENPF,* which is associated with the kinetochores during mitosis (Figure 2DD).^40^ This cluster consists of cells in a transitory cell cycle state that is unique to dividing cells. *SMC2* is expressed broadly in the undifferentiated epidermis at the base of the embryonic feather germ, but becomes more restricted towards the basal layer of epidermis once barb ridges form and epidermal feather cell types differentiate (Figure S2). *SMC2* expression identifies active areas of cell division but does not contribute to feather cell type identification (Figure S2).

#### Sheath Cell Types

The feather sheath is the deciduous outer layer of the developing feather germ that flakes apart and falls off as the feather matures. One large, complex cluster of epidermal cells (Figure 2A) consists of cells that strongly express *KRT18*, *NEBL*, *ALCAM*, and *CDH2* (Figure 2P, DD). *KRT18* is expressed in single layered epithelia, including human periderm.^41,42^ In situ hybridization shows that *KRT18* is expressed in the periderm of the embryonic chick skin between developing feather germs, and in the outermost layer of the sheath (Figure 2B, S4A). A single cluster of *KRT18* expressing periderm cells is present in our analyses of plucked feather samples (Figure 2A), but the feather and skin sample contains two *KRT18* expressing clusters (Figure S1A, L). One includes far fewer cells than the other. The larger of these two clusters (Figure S1A) also expresses *PSCA* (Figure S1M) which is shared by the periderm sheath cells identified from the plucked feather samples (Figure 2Q), and thus corresponds to the periderm sheath of the feather germ. The smaller cluster (Figure S1) corresponds to the interfollicular periderm present in this sample (Table 1).

A heterogeneous group of three epidermal cell clusters (Figure 2A) share expression of *ABCA12*, *DOCK9*, *GSN*, and *EZR* (Figures 2R and 2DD). *ABCA12*, a transmembrane lipid export protein,^43^ is expressed in the basal sheath cells in ramogenic stage feathers, and we identified these cell clusters as components of the feather sheath (Figure 2D, S4B). In Barbule Initiation Stage feathers, *ABCA12* is also expressed in the developing barbule plates, however we can distinguish these clusters from sheath cells using other marker genes (see below) (Figure 2E, DD). Although these three clusters represent a sheath cell type, there is some overlap in gene expression with the periderm sheath, such as *ANXA1*, *DOCK9*, and *EZR* (Figure 2S, DD). One of these three clusters of sheath cells (Figure 2A) shares expression of *ANXA1*, a phospholipid dependent membrane bound protein involved in membrane cross linking,^44^ with sheath periderm (Figure 2C, S, DD). *ANXA1* is expressed broadly in the sheath at all stages of feather germ development (Figure 2C). Although our analysis identified three sheath cell clusters, we are unable to distinguish these expression states anatomically and they may represent transitional maturation states.

#### Barb Ridge Cell Types

Barb ridge cells develop into the ramus and barbules that make up the natal down feather.^6^ A cluster of cells (Figure 2A) differentially expresses *NRCAM*, *AKAP12*, and *DIPK2A*. *NRCAM* (Figure 2T, DD) localizes to the undifferentiated barb ridge cells located at the base of the feather germ (Figure 2K). These cells appear to be a transitory cell state of undifferentiated barb ridge cells that occur after the development of barb ridges but prior to the differentiation of barb ridge cells into feather cells types. Thus, these cells are likely to be a developmental intermediate state within the barb ridge.

A cluster of cells (Figure 2A) were identified by expression of *AXL*, *NFKBIZ*, *KITLG*, and *FAM129A* (Figure 2U, DD). AXL is a receptor tyrosine kinase involved in signal transduction from the extracellular space to the cytoplasm.^45,46^ These *AXL*-expressing cells are anatomically located in the developing barbule plates in Barbule Initiation Stage feather cells (Figure 2G). Within the barbule plate, *AXL* expression starts superficially (i.e., peripherally) and expands basally (i.e., internally) as barbule development proceeds, congruent with the superficial-to-basal maturation gradient of the epidermis (see Supplementary Information). *AXL* expression (Figure 2G) replaces *NRCAM* expression (Figure 2K, L) as cells differentiate from transitional undifferentiated barb ridge cells into barbule plate cells.

A distinct cluster of cells (Figure 2B) that express *AXL*, *ESRRB*, *PLEKHA6*, *MCF2*, and *CRACD,* is only identifiable in later Ramus Development stage samples (Figure 2U, DD). In situ hybridization for *AXL* identified this cluster as cortical cells of the barb rami (Figure 2L). Thus, as barb ridge development proceeds, *AXL* expression advances from the barbule plate cells in Barbule Initiation Stage to the cortical cells of the barb ramus cells in during the Ramus Development stage (Figures 2G, L, and S3A-B).

A small, but highly distinct cluster (Figure 2A) of epidermal cells at the Ramus Development stage strongly express *CHL1*, *THSD7B*, *HEPHL1*, and *TSHZ1* (Figure 2V, DD). *CHL1* is also expressed in the undifferentiated epidermal cells (Figure S3A-B). The appearance of this cluster of cells corresponds with the development of medullary ramus cells, which are absent in the earlier Barbule Initiation Stage samples (Table 1). In situ stains confirm distinct *CHL1* expression in the medullary cells of the ramus during the Ramus Development stage (Figure 2M).

One large cluster of β-keratin expressing cells (Figure 2A) occurs only in the ramus initiation stage samples (Figures 2DD). These cells show lower RNA diversity than other epidermal clusters. We hypothesize that this cluster contains cells of multiple cell types that are convergently similar through the process of final keratinization and cell death.

#### Basal Epithelium

The basal epithelium of the feather germ generates two cells types that function critically in feather morphogenesis, but do not become enduring components of the mature feather. One large cluster (Figure 2A) of *SEMA5A*, *CADM1*, and *SDK1* expressing epidermal cells (Figures 2W-Y) contains two subclusters with distinct transcriptomic profiles.

One subcluster (Figure 2A) strongly expresses Sonic Hedgehog (*SHH*) and Bone Morphogenetic Protein 2 (*BMP2*) (Figure 2Z-AA). As expected from previous research,^21^ these cells localize to the marginal plate epithelium (Figure 2H-I). *SHH* is expressed broadly in the entire marginal plate, and *BMP2* expression is restricted to the apical folds of the marginal plate (Figure 2H-I).^21^ Later, during the Ramus Development Stage (See Supplement), *BMP2* is expressed more broadly within the epidermal feather cells (Figure S4C).

The second of subcluster of *SEMA5A*, *CADM1*, and *SDK1* expressing cells (Figure 2A) lacked *BMP2* and *SHH* expression, but had distinct expression of latent transforming growth factor beta binding protein 1 (*LTBP1*), which was absent in the marginal plate cells, as well as *SLC6A15*, *SLC7A11*, and *SLC1A4* in the solute carrier family (Figures 2BB and 2DD). *LTBP1* is involved in paracrine signal transduction,^47^ which implies a novel, and previously unanticipated role for the barb ridge basal epithelium in barb ridge, ramus, and barbule morphogenesis. We hypothesize that this cluster corresponds to the unfolded basal epithelium of the barb ridges. However, in situ staining for *LTBP1* was broadly distributed across all cell types (Figure S4D), and more work remains to distinguish the barb ridge basal epithelium cells from the marginal plate and the basal epithelium prior to barb ridge formation.

#### Axial Plate

The axial plate consists of a thin row of cells that lie between the barbule plates early in barbule development. These cells do not constitute a part of the mature feather. A distinct cluster of cells identified in analyses of the plucked Barbule Initiation Stage feathers (Table 1) expresses Semaphorin 3C (*SEMA3C*), which is involved in signal receptor activity. In situ hybridization documents *SEMA3C* expression in the axial plate cells that lie between the developing barbule plates in the Barbule Initiation Stage samples (Figure 2J). However, we were unable to reliably identify a distinct cell cluster corresponding to the axial plate in our aggregate analyses based on all plucked feather samples (Figure 2CC). This result may be due to the fact that axial plate cells are very limited in number and temporally ephemeral. However, this finding is the first identification of any distinct gene transcription state or paracrine signaling function for axial plate cells, and it confirms a distinct developmental function for this anatomically identified cell type.^6^

### RNA Velocity

RNA velocity analyses are used to infer a high dimensional vector field estimating the future transcriptional state of individual cells.^48^ We used RNA velocity to describe cellular differentiation in feather keratinocytes and inferred a logical development of each of the cell types identified by our single cell data.

Our undifferentiated feather epidermis appears to be the developmental precursor to all of the feather cell types except the sheath periderm (Figure 3). In general, vectors flow in three directions from the undifferentiated feather epidermis cluster: the basal epithelium clusters, the sheath clusters, and the barb ridge clusters. Some cells in this undifferentiated epidermal cell cluster had vectors pointing toward the cell cycle cluster, indicating that this cluster is actively dividing and differentiating.

The sheath clusters show long parallel arrows at the periphery, implying rapid differentiation and larger change in gene expression^49^ (However, vector length is confounded by the non-linear dimensional reduction used in UMAP). The direction of the arrows implies that sheath cells develop from the undifferentiated epidermis. We also see a mix of sheath cells undergoing differentiation (long parallel arrows), and differentiated sheath cells maintaining homeostasis prior to cell death (shorter arrows with more mixed directions). The periderm sheath appears on its own with no obvious developmental precursor. This supports conclusion that these cells develop from the periderm layer of the embryonic epidermis that precedes feather development.

The basal epithelium clusters differentiate directly from the undifferentiated feather epidermis cluster. The marginal plate epithelium appears to be a precursor to the barb ridge basal epithelium, which is congruent with previous observations^21^ of new barb ridge formation (see Discussion).

Another trajectory flows from the undifferentiated feather epidermis to the barb ridge cell type clusters. These cells first develop into the undifferentiated barb ridge cells marked by *NRCAM*. This cluster is an intermediate cell state and most of the arrows move through this cluster indicating that this cluster is temporary.

The undifferentiated barb ridge cells then differentiate into the barbule plate and ramus cells, although the actual ordering of this process is confounded by noise in the RNA velocity. The barbule plate, ramus cortex, and ramus medulla each have arrows pointing in new directions instead of the direction of any other cluster, indicating that these three cell types are terminally differentiated.

The late keratinocyte cluster, which we hypothesize consists of dead or dying cells, appears in the flow plot to be a source sheath, ramus cortex, and barbule plate cells (Figure 3). However, because cells of this cluster have lower RNA counts and consists mostly of beta keratin from feather cells that are keratinizing and dying, we expect that this is locally perturbing the RNA velocity analysis.

### Cell Type Evolution

Using the averaged expression values for each cluster for transcription factor genes only, we inferred a neighbor-joining cell type tree hypothesized to represent evolutionary relationships among the differentiated epidermal cell types of the developing feather germ (Figure 4).^50,51^ We excluded most of our hypothesized transitory cell states and undifferentiated clusters because they show intermediate gene expression states. We rooted this tree with the sheath periderm cells, which we hypothesize to be the earliest, evolutionarily diverging epidermal cell type still present in feathers, and shared with embryonic reptile skin prior to the origin of feathers.

We find that the sheath cell clusters (sheath 1, 2, and 3) form a monophyletic group, supporting our identification of these clusters as a common cell type containing three subclusters. However, the barb ridge cell types do not appear monophyletic on this tree. Rather, the barbule plate cells are placed as sister to the non-periderm sheath cells. A second clade contains the marginal plate, barb ridge basal epithelium, ramus cortex, and ramus medulla cell types, respectively.

This result suggests that there is a broad evolutionary divide between the basal and superficial epidermal cell types. The placement of the barbule plate cells with the sheath may reflect some overall gene expression signature of their proximate, superficial position next to the sheath. The two ramus cell types – cortical and medullary cells – form a monophyletic group, indicating that these two cell types evolved from a ramus specific cell type ancestor. Lastly, the very long branch of the ramus medulla cell type (Figure 4) indicates that this cell type is highly diverged transcriptomically from the ramus cortex despite their shared common ancestor.

## Discussion

Single cell transcriptomics of developing embryonic chick feathers confirms the existence of highly distinct cell expression clusters with diagnostic cell transcriptomic profiles. Specifically, we identified eight cell types that were previously characterized as having distinct morphologies and developmental histories by Lucas and Stettenheim (1972; Figure 241).^6^ Four other of morphological feather cell types proposed by Lucas and Stettenheim are not present in our data because embryonic chick feathers lack a rachis, pulp caps, and differentiated proximal and distal barbules. In addition, we found that the basal epithelium of the barb ridges is likely to be a distinct cell type from both the marginal plate epithelium and the undifferentiated basal epithelium prior to barb ridge formation. These cells are likely to play an important role in the development of pulp caps in feathers generated by molt in older birds. Likewise, we identified the periderm component of the embryonic feather sheath, which Lucas and Stettenheim did not recognize.

We confirm the results of Harris et al.^21,22^ that *SHH* is expressed broadly in the marginal plate epithelium during the development of barb ridges and barb cells types, while *BMP2* expression is restricted to the peripheral bend in the marginal plate (Figure 2H-I). We also identify the basal barb ridge epithelium as a closely related set of cells that lack SHH and BMP2, instead expressing *LTBP1*, *SLC6A15*, *SLC7A11*, and *SLC1A4* (Figures 2BB and 2DD). These results suggest that the marginal plate and basal barb ridge epithelium function together in both barb ridge formation and in the establishment of a superficial-basal signaling gradient that provides local spatial orientation information necessary for the morphogenesis and hierarchical organization of barbule and ramus cells.^22^

We also identified the first known molecular marker for axial plate cells: *SEMAC3C* (Figure 1J). *SEMAC3C* is an extracellular paracrine signaling protein and it indicates that axial plate cells function in barbule plate morphogenesis, although further work is necessary to investigate axial plate cell signaling, and establish the developmental function played by *SEMA3C* in these cells.

### Feather Cell Type Development

Our RNA velocity analysis provides a new perspective on the development of feathers through the derivation of their distinct cell types. In general, we find two distinct developmental trajectories for cells that do not contribute to the final mature feather morphology, and one trajectory for the structures that comprise the feather (Figure 3).

The basal epidermis trajectory branches off of the undifferentiated cells and towards the marginal plate epithelium, and onward to the marginal plate to the barb ridge basal epithelium, implying that the marginal plate develops prior to the barb ridge basal epithelium (Figure 3). This conclusion is supported by previous observations that barb ridges are developmentally initiated with a infolding of the peripheral bend of marginal plate between barb ridges.^22^ This new marginal plate expands before the basal epithelium of the barb ridge develops. Thus, the marginal plate is functionally required prior to the development of a distinct barb ridge basal epithelium.

The sheath cell type also appears to develop directly from undifferentiated epidermal cells (Figure 3). The differentiation of the sheath cells occurs early in feather development, and is stable throughout sheath keratinization. The periderm is already differentiated at the developmental stage of our sampling, so we are unable to map its developmental trajectory.

Another developmental pathway takes undifferentiated epidermal cells through a barb ridge-specific cell state (Figure 3). Although these cells differentially express *NRCAM*, it is transitory developmental state, which is later superseded by barbule, ramus cortical, and medullary cell identities (Figures 2K-M, 3, S3). The ramus cortex, ramus medullary cells, and barbule plate cell types each have vectors leading away from a central point, implying that these are each distinct differentiation pathways. Unlike most other animal cell types, which are maintained through time, feather cell types develop their identities, produce keratin, solidify, and die as they form the structure of the feather. Therefore, it is unsurprising that each of these cell types has arrows pointing away from the center of the cluster, but not towards any other cell type. This indicates a distinct developmental identity that will eventually keratinize and die.

### Cell type evolution

The developmental model of feather evolution^3^ proposed a hierarchical model of the origin and diversification of the feather morphology based on developmental data. The hypothesis that feathers evolved through a series of innovations in developmental mechanism also makes explicit predictions about the evolutionary sequence of origins of feather cell types. By comparing our results to the predictions of the developmental model, we gain further insight about the history of feather evolution.

Prior to the origin of feathers, the embryonic epidermis was ancestrally differentiated into stratified layers – periderm, subperiderm, alpha and beta stratum, and the stratum basale.^52–54^ Primitive scale placode cell types were subsequently gave rise to the cells of feather placodes.^55^ Likewise, various differentiated scale cells types were already present at the onset of feather evolution. Our cell type tree partially reflects this ancestral stratification of epidermal layers; the periderm, the outer epidermal cell types, and basal epidermal cell types all form clades, and the intermediate layer cell types are nearly monophyletic (Figure 4).

Stage I feathers in the developmental model^3^ consist of a tube of epidermis with a central dermal core that morphologically resembles the hollow calamus of a feather (but with unknown dimensions or mechanical properties). At this stage, the feather would have evolved solidly keratinized cells, that are probably similar to cortical cells of the barb rami or calamus. It is unknown specifically when the feather follicle or periodic molt evolved, but it is possible that these could have evolved at this stage. Furthermore, at Stage I, the dermal pulp would have likely evolved a proliferative cell type found at the base of the feather germ, and more connective tissue rich dermal pulp cells with the feather germ. The origin of pulp caps likely coincided with the evolution of molt in Stage I or with Stage II. Likewise, calamus could have evolved in Stage I feathers, but would have been necessary for the evolution of molt.

Stage II feathers consist of unbranched barbs, a deciduous sheath, a tubular calamus, and almost certainly iterative molt. The evolution of the Stage II feathers required the origin of several, novel cell types. A sheath cell type evolved from a combination of periderm and the outer epidermal layer. The evolution of barbs required the origin of solidly keratinized cells stratum intermedium, and marginal plate epithelium and barb ridge basal epithelium from the stratum basale. The origin of the differentiated medullary ramus cell type could have evolved at the same time as the first barbs, or later in response to selection on ramus mechanical function. Regardless, ramus medullary cells likely evolved before the evolution of doubly branched feather structure (Stage IIIa+b; see below).

Our phylogeny shows that the periderm and sheath are not sister to one another, suggesting that the sheath evolved from the subperiderm and the stratum intermedium (Figure 10). The sheath periderm and the other sheet cell types would have evolved with the origin of Stage II feathers. Interestingly, the embryonic feather sheath seems to be a composite structure of cells from different origins. However, because the periderm is only present in embryonic skin, all feather sheaths grown through molt later in life are composed of subperiderm and outer epidermal layer cells. Among the other barb ridge cells, we find that the marginal plate diverges before the barb ridge basal epithelium, matching the prediction of the developmental model.

Stage IIIa feathers consist of a rachis and multiple barbs without barbules, which required the origin of a new barb ridge locus, helical growth of barb ridges, and dorsal fusion of barb ridges to form the rachis, initially, and to fuse to the rachis thereafter. None of these novelties require new feather cell types, but involve further control of barb ridge morphogenesis ^21,22^. Although the rachis is absent from chick natal down and is not analyzed here, the rachis is predicted to consist of identical cell types to the ramus. We hypothesize that the distinct identity of the rachis is a product of barb ridge morphogenesis, not of the novel cell types within it.

Stage IIIb feathers consist of tuft of rami with multiple undifferentiated barbules. This morphology required the origin of paired barbule plates from the intermediate layer, and a row of axial plate cells between them. The first barbule cells would have been simple, strap-shaped “basal” cells. Stage IIIa+b feathers consist of a rachis with multiple barbs and barbules. It could have evolved by either a->b, or b->a.^3^

Our model places cortical and medullary cells as a cell type clade, supporting the idea that these differentiated cell types evolved from a common ancestral cell type that been present in Stage II feathers (Figure 4). Interestingly, the long branch leading to ramus medullary cells indicates a large amount of evolutionary change in gene regulation from a solidly keratinized, cortex-like ancestral state. The large, air-filled medullary cells are unique among vertebrate integumentary appendages, and maybe the most distinct feather cell type innovation.

The placement of the barbule cell type as more closely related to sheath than to barb ridge cell types conflicts with the devo-evo model’s cell type predictions. This inferred relationship could reflect the spatial position of the barbule plates as adjacent to the sheath and superficial to the ramus. However, it is also possible that the barbule cells have converged in some developmental or keratinization mechanisms on sheath cells.

Stage IV feathers consist of a coherent, pennaceous vane with differentiated proximal and distal barbules with hooklets and grooves, respectively. These morphologies involve the evolution of barbicels, or thin, keratinized extensions of barbule cell shape. Because barbicels are not present in chick natal down, we do not yet have data relevant to these microstructural novelties of the barbule cell type.

In conclusion, this transcriptomic investigation of developing chick feathers supports the existence of numerous cell types previously predicted by cellular morphology, and is substantially congruent with prediction of cell type evolutionary sequence derived from the developmental model of feather evolution.^3^ Our results suggest an expanded model of feather evolution based on a stepwise addition of specialized cell types responsible for the key morphological and developmental innovations, leading to the complex morphology and diversity of extant feathers. These novel cell types include those that contribute to the morphology of the mature feather, and those that function exclusively during the development of the feather.

## Supporting information

Supplemental Background

## Acknowledgements

We would like to thank all the members of the Prum Lab for their feedback and comments on this manuscript. Jamie Maziarz helped troubleshoot protocols. Daniel Stadtmauer helped with experimental methods and data analysis. We would also like to thank the UCONN Poultry Farm for providing fertilized eggs. We would also like to thank the Yale Center for Genome Analysis for sequencing consultations and protocol advice. Research reported in this publication was supported by the National Institute of General Medical Sciences of the National Institutes of Health under Award Number 1S10OD030363-01A1. This work is supported by the Yale University W.R. Coe Fund and the NIH Training Program in Genetics number 5T32GM007499-43.

## Author Contributions

C.L. and R.O.P. conceived of the project. C.L., R.O.P., and G.W. designed experiments. C.L. optimized protocols, conducted experiments, and analyzed data. C.L. wrote the first draft and R.O.P., G.P., and C.L. contributed to editing and refining.

## Supplemental Information Index

Document S1. Table S1. Figures S1-S4

## Methods

### Resource Availability

#### Lead Contact

Further information and requests for resources and reagents should be directed to and will be fulfilled by the lead contact, Cody Limber.

#### Materials availability

Probe accession numbers can be found in the resources table.

#### Data and code availability

Single-cell RNA-seq data have been deposited at GEO^56^ at GEO: GSE275984 and are publicly available as of the date of publication

All original code has been deposited at Zenodo at 10.5281/zenodo.13685474 and is publicly available as of the date of publication.

### Experimental model and subject details

White leghorn chicken (*Gallus gallus*) fertilized eggs were sourced from the University of Connecticut Poultry Farm (Storrs, Connecticut). Eggs were kept at room temperature before being placed in an incubator and kept at 37° C with a humidity of 40-60%. Eggs were automatically rotated every two hours. All embryos were staged according to Hamburger– Hamilton stages (1951).^57^

### Method Details

#### Cell and nuclear dissociation for scRNA-seq

We dissected H&H stage 38, 39, and 40 embryos^57^ in PBS on ice (see Table 1). For our sample containing skin and feathers, we dissected a section of skin from the dorsal feather tract peeled off most of the dermis leaving primarily epidermis and embryonic feathers. For feather only samples, we plucked feathers from a section of the dorsal feather tract just below the neck of the embryo. Dissected samples were placed in PBS on ice and kept for less than 15 minutes before proceeding to single cell or single nuclear isolation.

For samples from stage 38 embryos, we gently chopped the tissue with a scalpel before incubating the samples in a solution of Pronase (1mg/1ml PBS) for 60 minutes at 37° C. The tissue was then gently passed 20 times though an 18 gauge needle. The resulting cellular suspension was filtered through a 70μm filter and then filtered twice through a 40μm filter to remove larger debris. We then spun down the cell suspension at 300 g for 5 minutes removing the supernatant and resuspended the pellet in calcium and magnesium free PBS containing 1% bovine serum albumin twice to wash the cells. We used Trypan blue to estimate cellular viability with only a ≥ 70% viability used for further processing. We also estimated the cellular concentration on a hemocytometer before diluting the suspension to 1000 cells/μL to submit for 10x library preparation.

For samples from stage 39 and 40 embryos, we resorted to nuclear isolation as single cell dissociation was impossible due to the keratinization of cells in these stages. Our protocol was adapted from Sarropoulos et al.^58^ Plucked feathers were placed in 2 mL of TST lysis buffer containing 1ml 2x Salt-Tris Buffer (see below for 1x buffer), 60 μL 1% Tween-20, 10 μL 2% bovine serum albumin, and 930 μL nuclease free H2O. We ground the tissue using a dounce homogenizer grinding 20 times with the loose pestle and 20 times with the tight pestle. This suspension was filtered through a 70μm filter and then filtered twice through a 40μm filter. Nuclei were pelleted at 500 g for 5 minutes before being washed with a Salt-Tris Buffer (146 mM NaCL, 1mM CaCL2, 21mM MgCl2, 20mM tris-HCl PH 7.5). The nuclei were then and resuspended in ice cold PBS without calcium or magnesium and containing 1% bovine serum albumin for submission for 10x library preparation.

#### Single-cell cDNA library preparation

Cells and nuclei were kept on ice until submission to the Yale Center for Genomic Analysis for library preparation using a 10X Genomics Chromium platform for Chromium Single Cell 3′ v.3 Library prep targeting 10,000 cells. The resultant cDNA libraries were sequenced on an Illumina NovaSeq apparatus.

#### RNA in situ hybridization

Tissue for RNA in situ hybridization was collected from H&H stage 38, 39, and 40 embryos^57^ and fixed in 10% PFA overnight. We then cleared the tissue in xylenes, dehydrated in ethanol, and then embedded them in paraffin under vacuum. After sectioning, we followed the protocol available from ACDBio RNAscope^59^ for the RNAscope 2.5 High Definition (HD)— BROWN Assay to stain for marker genes of interest. Probe information is available in Table S1.

#### Microscopy

Images were also acquired on a Leica THUNDER Imager 3D Tissue. The microscope was controlled by the Leica LAS X software and we stitched images from a single plane together using Leica Navigator in Leica LAS X. Final images were brightened, color balanced and arranged in Photoshop 2024 (version 25.6).

Images were also acquired using an Olympus BH-51 compound microscope outfitted with a Teledyne Lumenera Infinity 3 digital camera. Positioning of the slide, focus, and image acquisition were controlled by software (Objective Imaging Surveyor version 9.4.0.5). Individual position images were stitched together on a single plane, and multiple planes rendered into an extended focus image using Helicon Focus Pro version 8.2.0. Final images were brightened, color balanced and arranged in Photoshop 2024 (version 25.6).

### Quantification and statistical analysis

#### Cluster Analysis

Sequencing data were demultiplexed and aligned to the Ensembl GRCg6a genome by 10x Genomics Cell Ranger version 7.0.0^60^ for generation of the gene by cell count matrices. Ambient RNA filtering was performed by SoupX version 1.6.2^61^ prior to standard filtering, normalization, and quality control steps which were performed using the Seurat (version 4.1.1^62^) software package in R, version 4.3.2. Data were filtered to exclude droplets with <1000 features/cell to exclude empty droplets. These data were then normalized to counts per 10,000 prior to integrating our plucked feather datasets using seurat anchoring for integration and batch correction.

We ran a principal component analysis and UMAP on 20 dimensions to visualize the data. We also ran FindNeighbors (dims 1:20) and FindClusters (resolution 0.5) to identify cell clusters. We filtered out two clusters, one containing primarily cells with a high proportion of mitochondrial counts indicating damaged cell membranes, and one containing markedly high features/cell indicating presence of doublets. We then reran UMAP to visualize the remaining cells. These cell clusters were used to identify top differentially expressed marker genes that we used for cell type identification. We also identified 12 clusters pertaining to the epidermal feather cell types which we subsetted and visualized using UMAP. Two of these clusters (ramus and marginal plate) were subsetted since we found our initial clustering was unable to capture the variation in expressed genes we observed within these clusters.

#### RNA Velocity

To determine cellular developmental trajectories, we performed RNA velocity analysis on our epidermal feather cell subset. We used Velocyto version 0.17.17^48^ to count spliced and unspliced RNAs and scVelo version 0.2.5^63^ in Python version 3.10.8 to compute velocities and visualize cell RNA velocity vectors.

#### Cell type tree

We calculated pairwise distances between averaged cluster expression of transcription factors. To create cell type trees, we used the neighbor joining package in Ape^64^ version 5.6-2 in R version 4.3.2.

## References

1. Shapiro, E., Biezuner, T., and Linnarsson, S. (2013). Single-cell sequencing-based technologies will revolutionize whole-organism science. Nature Reviews Genetics 14, 618.

2. Wagner, G.P. (2014). Homology, genes, and evolutionary innovation (princeton university press).

3. Prum, R.O. (1999). Development and evolutionary origin of feathers. Journal of Experimental Zoology 285, 291–306.

4. Wagner, G.P., Chiu, C.-H., and Laubichler, M. (2000). Developmental evolution as a mechanistic science: the inference from developmental mechanisms to evolutionary processes. American Zoologist 40, 819–831.

5. Arendt, D., Musser, J.M., Baker, C.V., Bergman, A., Cepko, C., Erwin, D.H., Pavlicev, M., Schlosser, G., Widder, S., and Laubichler, M.D. (2016). The origin and evolution of cell types. Nature Reviews Genetics 17, 744.

6. Lucas, A.M., and Stettenheim, P. (1972). Avian anatomy: integument (US Agricultural Research Service).

7. Clark, C.J., and Feo, T.J. (2008). The Anna’s hummingbird chirps with its tail: a new mechanism of sonation in birds. Proceedings of the Royal Society B: Biological Sciences 275, 955–962.

8. Wolf, B.O., and Walsberg, G.E. (2000). The role of the plumage in heat transfer processes of birds. American Zoologist 40, 575–584.

9. Rijke, A.M. (1968). The water repellency and feather structure of cormorants, Phalacrocoracidae. Journal of Experimental Biology 48, 185–189.

10. Terrill, R.S., and Shultz, A.J. (2023). Feather function and the evolution of birds. Biological Reviews 98, 540–566.

11. Hill, G.E., and McGraw, K.J. (2006). Bird coloration (Harvard University Press).

12. Baker, R.R., and Parker, G.A. (1979). The evolution of bird coloration. Philosophical Transactions of the Royal Society of London. B, Biological Sciences 287, 63–130.

13. Clark, C.J., Elias, D.O., and Prum, R.O. (2013). Hummingbird feather sounds are produced by aeroelastic flutter, not vortex-induced vibration. Journal of Experimental Biology 216, 3395–3403.

14. Stoddard, M.C., and Prum, R.O. (2008). Evolution of avian plumage color in a tetrahedral color space: a phylogenetic analysis of new world buntings. The American Naturalist 171, 755–776.

15. Stettenheim, P. (1976). Structural adaptations in feathers. (Australian Academy of Science Canberra, Australia), pp. 385–401.

16. Chen, C.-F., Foley, J., Tang, P.-C., Li, A., Jiang, T.X., Wu, P., Widelitz, R.B., and Chuong, C.M. (2015). Development, regeneration, and evolution of feathers. Annu. Rev. Anim. Biosci. 3, 169–195.

17. Strong, R.M. (1902). The development of the definitive feather (Bulletin of the Museum of Comparative Zoology at Harvard College).

18. Chodankar, R., Chang, C.-H., Yue, Z., Jiang, T.-X., Suksaweang, S., Burrus, L.W., Chuong, C.-M., and Widelitz, R.B. (2003). Shift of localized growth zones contributes to skin appendage morphogenesis: role of the Wnt/β-catenin pathway. Journal of investigative dermatology 120, 20–26.

19. Chu, Q., Cai, L., Fu, Y., Chen, X., Yan, Z., Lin, X., Zhou, G., Han, H., Widelitz, R.B., and Chuong, C.-m. (2014). Dkk2/Frzb in the dermal papillae regulates feather regeneration. Developmental biology 387, 167–178.

20. Yue, Z., Jiang, T.-X., Widelitz, R.B., and Chuong, C.-M. (2005). Mapping stem cell activities in the feather follicle. Nature 438, 1026–1029.

21. Harris, M.P., Fallon, J.F., and Prum, R.O. (2002). Shh-Bmp2 signaling module and the evolutionary origin and diversification of feathers. Journal of Experimental Zoology 294, 160–176.

22. Harris, M.P., Williamson, S., Fallon, J.F., Meinhardt, H., and Prum, R.O. (2005). Molecular evidence for an activator–inhibitor mechanism in development of embryonic feather branching. Proceedings of the National Academy of Sciences 102, 11734–11739.

23. Wong, H.H., Seet, S.H., Bascom, C.C., Isfort, R.J., and Bard, F. (2020). Red-COLA1: a human fibroblast reporter cell line for type I collagen transcription. Scientific reports 10, 19723.

24. Liu, X., Tang, H., Wang, Z., Huang, C., Zhang, Z., She, X., Wu, M., and Li, G. (2012). F10 gene hypomethylation, a putative biomarker for glioma prognosis. Journal of neuro-oncology 107, 479–485.

25. Urano, T., Narusawa, K.i., Shiraki, M., Sasaki, N., Hosoi, T., Ouchi, Y., Nakamura, T., and Inoue, S. (2011). Single-nucleotide polymorphism in the hyaluronan and proteoglycan link protein 1 (HAPLN1) gene is associated with spinal osteophyte formation and disc degeneration in Japanese women. European Spine Journal 20, 572–577.

26. Kou, I., and Ikegawa, S. (2004). SOX9-dependent and-independent transcriptional regulation of human cartilage link protein. Journal of Biological Chemistry 279, 50942–50948.

27. Lip, G.Y., and Blann, A. (1997). von Willebrand factor: a marker of endothelial dysfunction in vascular disorders? Cardiovascular research 34, 255–265.

28. Breier, G., Breviario, F., Caveda, L., Berthier, R., Schnurch, H., Gotsch, U., Vestweber, D., Risau, W., and Dejana, E. (1996). Molecular cloning and expression of murine vascular endothelial-cadherin in early stage development of cardiovascular system.

29. Alarcon-Martinez, L., Yilmaz-Ozcan, S., Yemisci, M., Schallek, J., Kılıç, K., Can, A., Di Polo, A., and Dalkara, T. (2018). Capillary pericytes express α-smooth muscle actin, which requires prevention of filamentous-actin depolymerization for detection. elife 7, e34861.

30. Rockey, D.C., Weymouth, N., and Shi, Z. (2013). Smooth muscle α actin (Acta2) and myofibroblast function during hepatic wound healing. PloS one 8, e77166.

31. Baek, S.-H., Maiorino, E., Kim, H., Glass, K., Raby, B.A., and Yuan, K. (2022). Single cell transcriptomic analysis reveals organ specific pericyte markers and identities. Frontiers in cardiovascular medicine 9, 876591.

32. Xia, M., Jiao, L., Wang, X.-H., Tong, M., Yao, M.-D., Li, X.-M., Yao, J., Li, D., Zhao, P.-Q., and Yan, B. (2023). Single-cell RNA sequencing reveals a unique pericyte type associated with capillary dysfunction. Theranostics 13, 2515.

33. Aalto, Y., El-Rifai, W., Vilpo, L., Ollila, J., Nagy, B., Vihinen, M., Vilpo, J., and Knuutila, S. (2001). Distinct gene expression profiling in chronic lymphocytic leukemia with 11q23 deletion. Leukemia 15, 1721–1728.

34. Strong, R.M. (1902). The development of color in the definitive feather. Science 15, 527–527.

35. D’Mello, S.A., Finlay, G.J., Baguley, B.C., and Askarian-Amiri, M.E. (2016). Signaling pathways in melanogenesis. International journal of molecular sciences 17, 1144.

36. Reemann, P., Reimann, E., Ilmjärv, S., Porosaar, O., Silm, H., Jaks, V., Vasar, E., Kingo, K., and Kõks, S. (2014). Melanocytes in the skin–comparative whole transcriptome analysis of main skin cell types. PLoS One 9, e115717.

37. Haake, A.R., König, G., and Sawyer, R.H. (1984). Avian feather development: relationships between morphogenesis and keratinization. Developmental biology 106, 406–413.

38. Strunnikov, A.V., Hogan, E., and Koshland, D. (1995). SMC2, a Saccharomyces cerevisiae gene essential for chromosome segregation and condensation, defines a subgroup within the SMC family. Genes & development 9, 587–599.

39. Kouprina, N., Pavlicek, A., Collins, N.K., Nakano, M., Noskov, V.N., Ohzeki, J.-I., Mochida, G.H., Risinger, J.I., Goldsmith, P., and Gunsior, M. (2005). The microcephaly ASPM gene is expressed in proliferating tissues and encodes for a mitotic spindle protein. Human molecular genetics 14, 2155–2165.

40. Liao, H., Winkfein, R., Mack, G., Rattner, J., and Yen, T. (1995). CENP-F is a protein of the nuclear matrix that assembles onto kinetochores at late G2 and is rapidly degraded after mitosis. The Journal of cell biology 130, 507–518.

41. Moll, R., Franke, W.W., Schiller, D.L., Geiger, B., and Krepler, R. (1982). The catalog of human cytokeratins: patterns of expression in normal epithelia, tumors and cultured cells. Cell 31, 11–24.

42. Dale, B.A., Holbrook, K.A., Kimball, J.R., Hoff, M., and Sun, T.-T. (1985). Expression of epidermal keratins and filaggrin during human fetal skin development. The Journal of cell biology 101, 1257–1269.

43. Akiyama, M. (2014). The roles of ABCA12 in epidermal lipid barrier formation and keratinocyte differentiation. Biochimica et Biophysica Acta (BBA)-Molecular and Cell Biology of Lipids 1841, 435–440.

44. Berg Klenow, M., Iversen, C., Wendelboe Lund, F., Mularski, A., Busk Heitmann, A.S., Dias, C., Nylandsted, J., and Simonsen, A.C. (2021). Annexins A1 and A2 Accumulate and Are Immobilized at Cross-Linked Membrane–Membrane Interfaces. Biochemistry 60, 1248–1259.

45. Lemke, G. (2013). Biology of the TAM receptors. Cold Spring Harbor perspectives in biology 5, a009076.

46. Dagamajalu, S., Rex, D., Palollathil, A., Shetty, R., Bhat, G., Cheung, L.W., and Prasad, T.K. (2021). A pathway map of AXL receptor-mediated signaling network. Journal of Cell Communication and Signaling 15, 143–148.

47. Altmann, C.R., Chang, C., Munoz-Sanjuan, I., Bell, E., Heke, M., Rifkin, D.B., and Brivanlou, A.H. (2002). The latent-TGFβ-binding-protein-1 (LTBP-1) is expressed in the organizer and regulates nodal and activin signaling. Developmental Biology 248, 118–127.

48. La Manno, G., Soldatov, R., Zeisel, A., Braun, E., Hochgerner, H., Petukhov, V., Lidschreiber, K., Kastriti, M.E., Lönnerberg, P., and Furlan, A. (2018). RNA velocity of single cells. Nature 560, 494–498.

49. Svensson, V., and Pachter, L. (2018). RNA velocity: molecular kinetics from single-cell RNA-Seq. Molecular cell 72, 7–9.

50. Arendt, D., Bertucci, P.Y., Achim, K., and Musser, J.M. (2019). Evolution of neuronal types and families. Current Opinion in Neurobiology 56, 144–152. 10.1016/j.conb.2019.01.022.

51. Kin, K., Nnamani, M.C., Lynch, V.J., Michaelides, E., and Wagner, G.P. (2015). Cell-type phylogenetics and the origin of endometrial stromal cells. Cell reports 10, 1398–1409.

52. Sawyer, R.H., and Knapp, L.W. (2003). Avian skin development and the evolutionary origin of feathers. Journal of Experimental Zoology Part B: Molecular and Developmental Evolution 298, 57–72.

53. Sawyer, R.H., Salvatore, B.A., Potylicki, T.T.F., French, J.O., Glenn, T.C., and Knapp, L.W. (2003). Origin of feathers: Feather beta (β) keratins are expressed in discrete epidermal cell populations of embryonic scutate scales. Journal of Experimental Zoology Part B: Molecular and Developmental Evolution 295, 12–24.

54. Sawyer, R.H., Rogers, L., Washington, L., Glenn, T.C., and Knapp, L.W. (2005). Evolutionary origin of the feather epidermis. Developmental dynamics: an official publication of the American Association of Anatomists 232, 256–267.

55. Musser, J.M., Wagner, G.P., Liang, C., Stabile, F.A., Cloutier, A., Baker, A.J., and Prum, R.O. (2018). Subdivision of ancestral scale genetic program underlies origin of feathers and avian scutate scales. bioRxiv, 377358.

56. Edgar, R., Domrachev, M., and Lash, A.E. (2002). Gene Expression Omnibus: NCBI gene expression and hybridization array data repository. Nucleic Acids Res 30, 207–210. 10.1093/nar/30.1.207.

57. Hamburger, V., and Hamilton, H.L. (1951). A series of normal stages in the development of the chick embryo. Journal of morphology 88, 49–92.

58. Sarropoulos, I., Sepp, M., Frömel, R., Leiss, K., Trost, N., Leushkin, E., Okonechnikov, K., Joshi, P., Giere, P., and Kutscher, L.M. (2021). Developmental and evolutionary dynamics of cis-regulatory elements in mouse cerebellar cells. Science 373, eabg4696.

59. Wang, F., Flanagan, J., Su, N., Wang, L.C., Bui, S., Nielson, A., Wu, X., Vo, H.T., Ma, X.J., and Luo, Y. (2012). RNAscope: a novel in situ RNA analysis platform for formalin-fixed, paraffin-embedded tissues. J Mol Diagn 14, 22–29. 10.1016/j.jmoldx.2011.08.002.

60. Zheng, G.X., Terry, J.M., Belgrader, P., Ryvkin, P., Bent, Z.W., Wilson, R., Ziraldo, S.B., Wheeler, T.D., McDermott, G.P., and Zhu, J. (2017). Massively parallel digital transcriptional profiling of single cells. Nature communications 8, 1–12.

61. Young, M.D., and Behjati, S. (2020). SoupX removes ambient RNA contamination from droplet-based single-cell RNA sequencing data. Gigascience 9, giaa151.

62. Hao, Y., Stuart, T., Kowalski, M.H., Choudhary, S., Hoffman, P., Hartman, A., Srivastava, A., Molla, G., Madad, S., and Fernandez-Granda, C. (2023). Dictionary learning for integrative, multimodal and scalable single-cell analysis. Nature Biotechnology, 1–12.

63. Bergen, V., Lange, M., Peidli, S., Wolf, F.A., and Theis, F.J. (2020). Generalizing RNA velocity to transient cell states through dynamical modeling. Nature biotechnology 38, 1408–1414.

64. Paradis, E., and Schliep, K. (2018). ape 5.0: an environment for modern phylogenetics and evolutionary analyses in R. Bioinformatics 35, 526–528. 10.1093/bioinformatics/bty633.

