## Supplemental Background for "Genetic Characterization of the Cell Types of in Developing Feathers, and the Evolution of Feather Complexity"

#### Feather Morphology

Branched feathers have a long central shaft, or rachis, that connects to the tubular calamus, or quill, at its base, which inserts into the skin. Branching off of the rachis are numerous barbs composed of a central ramus (pl. rami) and many smaller, secondary branches called barbules. Barb rami are multicellular in cross-section and typically consist of a solid outer cortex of completely keratinized cells and an inner medullary layer of hollow, air-filled cells. Barbules consist of a chain of single cells fused together, each of which is solidly keratinized (like the cortex of the barb ramus).^1^

The diversity of feather structures results from variation in the shapes and interactions of these hierarchically modular parts. Microscopic morphological differentiation between the hooked distal barbules and the grooved proximal barbules creates the opportunity for a Velcro-like zippering interaction that forms the closed pennaceous vane of a contour or flight feather. In contrast, the long, thread-like barbules and rami of downy feathers create puffy, space-filling structures of downy, or plumulaceous, feathers. Here, we investigate the development of chick natal down feathers, which consist of a tuft of bars with simple, undifferentiated barbules and no rachis.

#### Feather Development

The development of the first feathers and their follicles begins in the embryo with a feather placode, which elongates into a feather germ, or short bud (a tube of epidermis surrounding dermal pulp at its center). The feather follicle is formed as the epidermis proliferates around the base of the short bud and invaginates into the dermis. As cells divide at the base of the feather follicle, they replace older cells and push them upwards. Thus, the youngest cells are at the base of the feather and the oldest cells are at the tip. The mature feather is formed from the epidermal layer of the feather germ, while the dermal pulp provides nutrition to the growing feather.

The outermost epidermal layer of the feather germ forms the highly keratinized, deciduous sheath that protects the feather germ during growth and falls apart when the feather matures. The intermediate layer of epidermis forms the barb ridges which make up the branches of the feather. During development, the intermediate layer becomes subdivided into a series of barb ridges that become the barbs and rachis of the feather. Within a barb ridge, the barbules develop from barbule plates, or paired rows of cells extending toward the periphery of the feather germ. The barbule plates are separated by a single layer of axial plate cells, which do not become part of the mature feather. The barb ramus develops from the inner-most cells of the barb ridge. The basal layer of epidermis, or basal epithelium, separates barb ridges from the dermal pulp. As barb ridges form, they develop folds in the basal, or marginal plate, epithelium that separate neighboring barb ridges from each other.^2^

Previous research defines cell types as populations of cells that maintain consistent expression of a gene coregulatory complex ^3^. This criterion cannot be directly applied to feathers, or other integumentary appendages, because all cells reach their mature functional state as they die. Specifically, maturing feathers produce beta-keratin that polymerizes into an insoluble, intracellular solid, beginning on the periphery of the cell membrane and cutting them off from nutrients. Feather cells specify their identities and grow into the required shapes prior to keratinization. Thus, there is a window during which the feather cell identity is specified as the feather cells conforms to its adult shape while the cell is still alive. However, cell type specific transcriptomes cannot be maintained throughout keratinization and apoptosis.

#### Feather Developmental Stages

A single, developing feather germ has three gradients of maturation: a superficial–basal gradient within the layered epidermis, an apical-basal gradient across the tubular feather germ, and an anterior-posterior gradient within a transverse section of the tubular feather germ (especially pronounced in rachidial feathers, but present in chick down as well).^1,2,4-7^ The first, superficial–basal maturation gradient is characteristic of all epidermis generally, but becomes reoriented into a peripheral-interior gradient as the tubular feather germ grows out of the surrounding epidermis.

The onset of embryonic feather development varies over the surface of the body; initiation of feather tract development begins along the spine and expands like a spatial wave over the body.^1^ As a result, developmental state of feather germ cells is correlated not only with the age of the embryo, but also the position of those feathers on the body of the bird and the position of those cells within those feathers. Thus, single feather germs may contain cells at multiple stages of feather cell development.

To clarify the stages of development of the feather cells used in this study, we use the following stages to characterize the most advanced developmental state present in the feather samples analyzed at different embryonic development (H&H) stages.^8^

*Placode Stage*: At the placode stage, the site of feather development consists of a thickened epidermis with an underlying condensation of dermal cells. Placodes begin to develop on the dorsal midline of the back around H&H stage 30. We did not include the feather placode stage in our analysis.^9^

*Short Bud Stage*: The earliest emerging tubular feathers consist of a differentiated sheath covering undifferentiated epidermal feather cells. These epidermal cells are morphologically homogenous and are arranged in a tube surrounding a dermal core. Short bud stages begin to appear on the dorsal midline at H&H stage 35.

*Ramogenic Stage*: Developing barb ridges initiate the expansion of the folds of marginal plate epithelium that separate undifferentiated epidermal cells into barb ridges.^2^ The ramogenic stage begins to appear in the dorsal feather tract at H&H stage 37+.

*Barbule Initiation Stage*: The first event in the differentiation of barb ridge cells is the development of the most superficial cells into two parallel barbule plates separated by a single row of axial plate cells. Within each barbule plate, the more basal cells become the base of the barbules that fuse to the ramus, and the more peripheral cells become the tips of the barbules. Within the barb ridge, the more basal cells that will develop into the barb ramus are still undifferentiated. Feathers in barbule initiation stage begin to appear on the dorsal midline at H&H stage 38.

*Ramus Development Stage*: First, the basal-most cells within the barb ridge begin to differentiate into the cortical cells that form the outer surface of the ramus. Then, spongy medullary cells at the core of the ramus develop.^10^ By this stage, the barbule cells are nearly fully keratinized, and the elongate barbules extend upward within the sheath towards the distal tip of the feather (Lucas and Stettenheim, 1972; Figure 239).^1^ The ramus cells are the last to complete keratinization. Feathers in ramus development stage begin to appear on the dorsal midline at H&H stage 39.

To control for variation in the timing of feather placode initiation across the body, we sampled only from the dorsal midline of the back, where feather development first begins (H&H stage 30). (Table 1).

#### Feather Evolution

Prum (1999) proposed a model of feather evolution based on the hierarchal modularity of feather development. This structuralist approach hypothesized that feathers evolved through a series of evolutionary innovations in feather development, based on the development and diversity of extant feathers. The model was subsequently supported by both fossil data ^11-14^ and molecular developmental data.^2,7^

The developmental model of feather evolution proposed that feathers originated as a simple tube of epidermis surrounding a core of dermis (Stage I). Subsequently, the evolution of the capacity to subdivide the tube into barb ridges resulted in a tuft-like feather with a sheath (Stage II). Feathers then evolved either a helical growth, a new barb ridge locus, and a rachis (Stage IIIa), or barbule plates producing paired branches on barb rami (Stage IIIb). The evolution of both the ramus and barbules (Stage IIIa+b), were necessary to create the first doubly branched feather, but the relative order of IIIa and IIIb is still unestablished. The evolution of the closed pennaceous vane (Stage IV) required the origin of the capacity to differentiate proximal and distal barbules, allowing them to attach and interlock with one another. Feathers could then be elaborated upon by displacement of the new barb locus causing an asymmetrical feather vane (Stage Va) or addition of a new cite of barb ridge addition resulting in an afterfeather (Stage Vb)^15^.

The developmental model also provides novel hypotheses about feather cell type evolution (see Discussion). Given the morphological diversity available in chick natal down, we test this hypothesis by creating a phylogeny of feather cell types based on single cell sequencing data.^16^

### Supplemental Background References

1.	Lucas, A.M., and Stettenheim, P. (1972). Avian anatomy: integument (US Agricultural Research Service).

2.	Harris, M.P., Fallon, J.F., and Prum, R.O. (2002). Shh‐Bmp2 signaling module and the evolutionary origin and diversification of feathers. Journal of Experimental Zoology *294*, 160-176.

3.	Arendt, D., Musser, J.M., Baker, C.V., Bergman, A., Cepko, C., Erwin, D.H., Pavlicev, M., Schlosser, G., Widder, S., and Laubichler, M.D. (2016). The origin and evolution of cell types. Nature Reviews Genetics *17*, 744.

4.	Haake, A.R., König, G., and Sawyer, R.H. (1984). Avian feather development: relationships between morphogenesis and keratinization. Developmental biology *106*, 406-413.

5.	Lin, J., and Yue, Z. (2018). Coupling of apical-basal polarity and planar cell polarity to interpret the Wnt signaling gradient in feather development. Development *145*, dev162792.

6.	Prum, R.O., and Dyck, J. (2003). A hierarchical model of plumage: morphology, development, and evolution. Journal of Experimental Zoology Part B: Molecular and Developmental Evolution *298*, 73-90.

7.	Harris, M.P., Williamson, S., Fallon, J.F., Meinhardt, H., and Prum, R.O. (2005). Molecular evidence for an activator–inhibitor mechanism in development of embryonic feather branching. Proceedings of the National Academy of Sciences *102*, 11734-11739.

8.	Hamburger, V., and Hamilton, H.L. (1951). A series of normal stages in the development of the chick embryo. Journal of morphology *88*, 49-92.

9.	Musser, J.M., Wagner, G.P., Liang, C., Stabile, F.A., Cloutier, A., Baker, A.J., and Prum, R.O. (2018). Subdivision of ancestral scale genetic program underlies origin of feathers and avian scutate scales. bioRxiv, 377358.

10.	Prum, R.O., Dufresne, E.R., Quinn, T., and Waters, K. (2009). Development of colour-producing β-keratin nanostructures in avian feather barbs. Journal of the Royal Society Interface *6*, S253-S265.

11.	Wu, P., Hou, L., Plikus, M., Hughes, M., Scehnet, J., Suksaweang, S., Widelitz, R.B., Jiang, T.-X., and Chuong, C.-M. (2004). Evo-Devo of amniote integuments and appendages. The International journal of developmental biology *48*, 249.

12.	Prum, R.O., and Brush, A.H. (2002). The evolutionary origin and diversification of feathers. The Quarterly review of biology *77*, 261-295.

13.	Chen, C.-F., Foley, J., Tang, P.-C., Li, A., Jiang, T.X., Wu, P., Widelitz, R.B., and Chuong, C.M. (2015). Development, regeneration, and evolution of feathers. Annu. Rev. Anim. Biosci. *3*, 169-195.

14.	Ji, Q., Norell, M.A., Gao, K.-Q., Ji, S.-A., and Ren, D. (2001). The distribution of integumentary structures in a feathered dinosaur. Nature *410*, 1084-1088.

15.	Prum, R.O. (1999). Development and evolutionary origin of feathers. Journal of Experimental Zoology *285*, 291-306.

16.	Arendt, D., Bertucci, P.Y., Achim, K., and Musser, J.M. (2019). Evolution of neuronal types and families. Current Opinion in Neurobiology *56*, 144-152. <https://doi.org/10.1016/j.conb.2019.01.022>.
