## Supplemental Tables and Figures for "Genetic Characterization of the Cell Types of in Developing Feathers, and the Evolution of Feather Complexity"

Table S1: Probe accession info for probes used in situ hybridizations in this study

| Probe Name | Catalog Number |
| --- | --- |
| RNAscope 2.5 LS Probe- Gg-CDH5 | CAT NO: 458221 |
| RNAscope 2.5 LS Probe- Gg-ACTA2 | CAT NO: 1169151-C1 |
| RNAscope 2.5 LS Probe- Gg-F10 | CAT NO: 1196381-C1 |
| RNAscope 2.5 LS Probe- Gg-HAPLN1 | CAT NO: 1196391-C1 |
| RNAscope 2.5 LS Probe- Gg-KRT18 | CAT NO: 1178441-C1 |
| RNAscope 2.5 LS Probe- Gg-ANXA1 | CAT NO: 458191 |
| RNAscope 2.5 LS Probe- Gg-ABCA12 | CAT NO: 1169131-C1 |
| RNAscope 2.5 LS Probe- Gg-CHL1 | CAT NO: 1169001-C1 |
| RNAscope 2.5 LS Probe- Gg-AXL | CAT NO: 1169021-C1 |
| RNAscope 2.5 LS Probe- Gg-SHH | CAT NO: 551581 |
| RNAscope 2.5 LS Probe- Gg-BMP2 | CAT NO: 560601 |
| RNAscope 2.5 LS Probe- Gg-SEMA3C | CAT NO: 1169011-C1 |
| RNAscope 2.5 LS Probe- Gg-NRCAM | CAT NO: 1169041-C1 |

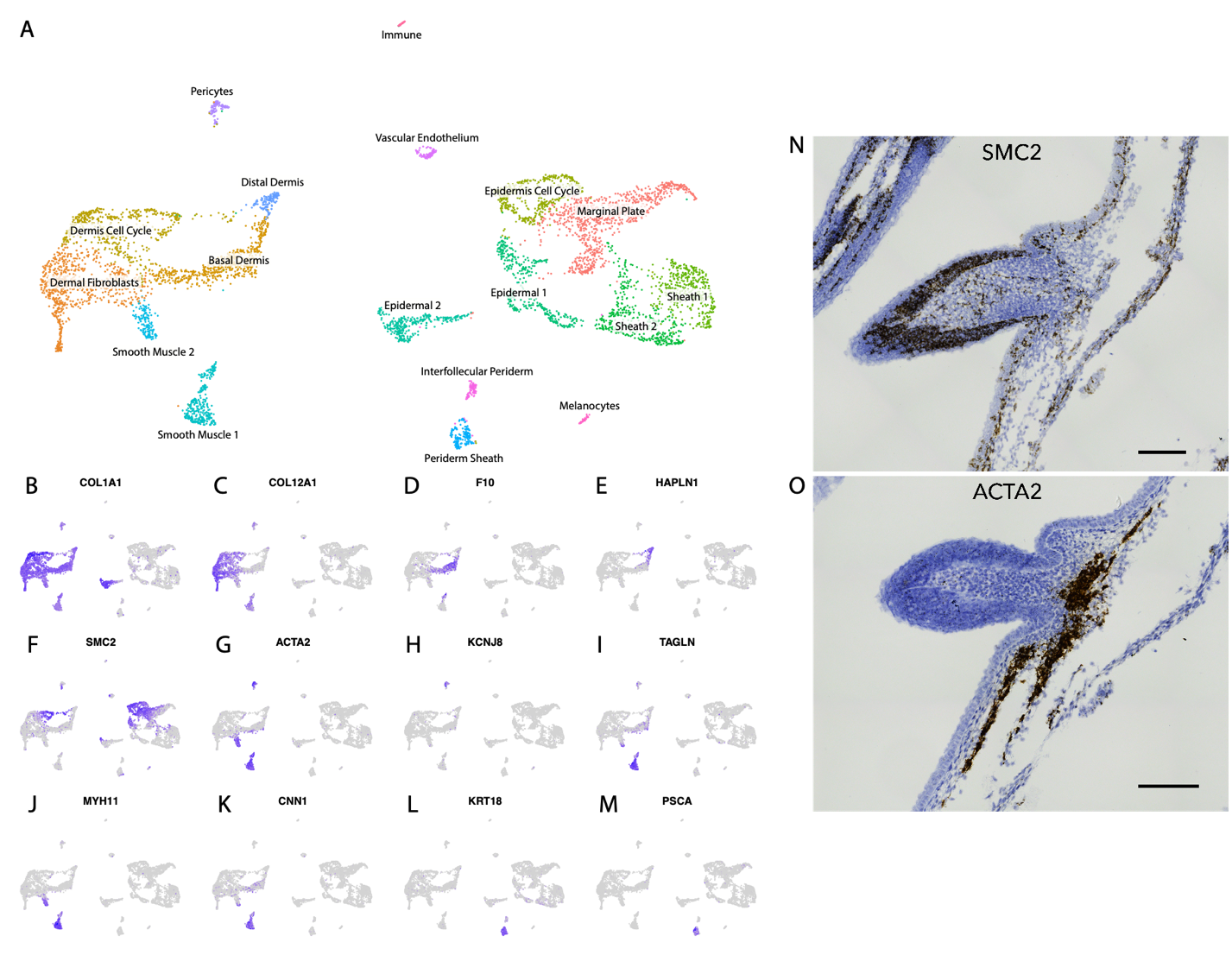

Figure S1: Analysis of non-feather skin cells from the feather and skin sample (Table 1), related to Figures 1 and 2

(A) UMAP plot of cells from the feather and skin sample colored and labeled by cell type identity.

(B-M) Feature plots highlighting expression levels for specific marker genes used in non-feather skin cell type cluster identification.

(N-O) In situ hybridization stained histological sections of genes used in non-feather skin cell type identification. Scale bars 0.1mm.

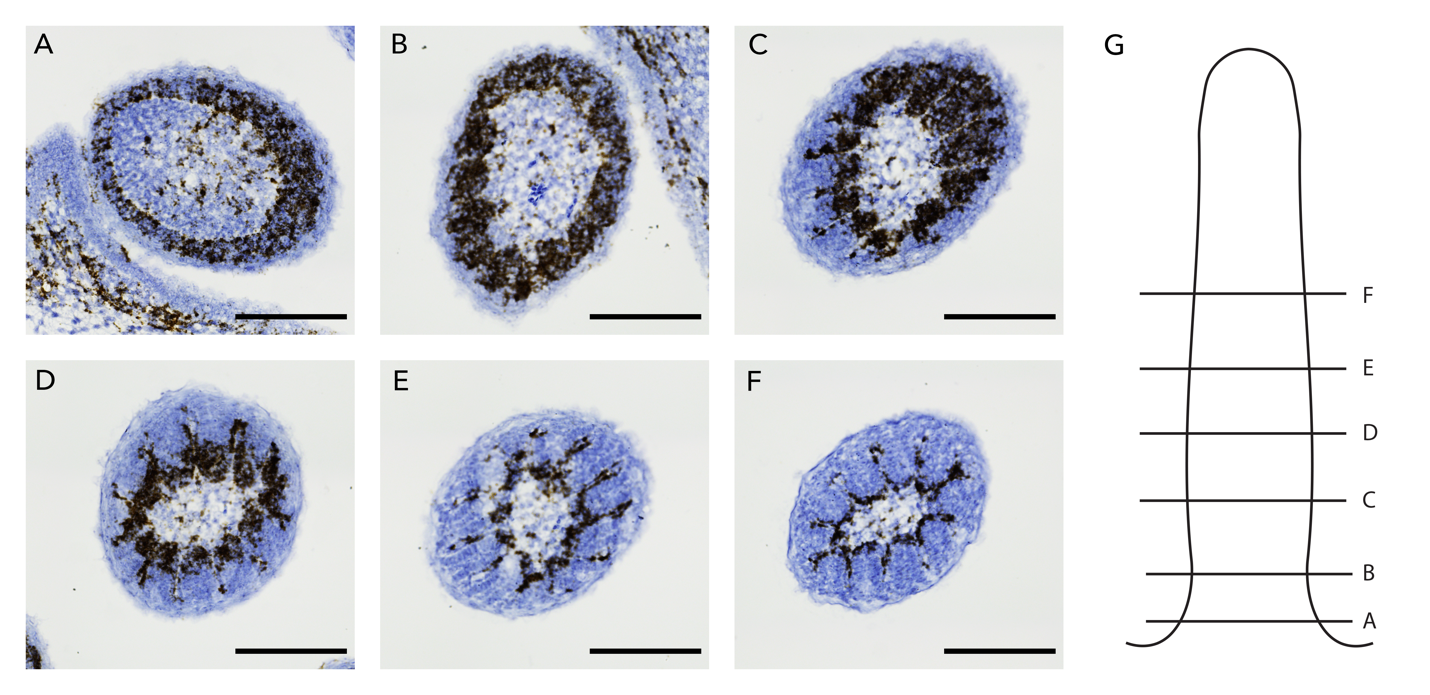

Figure S2: Changes in SMC2 expression over time, related to Figure 2

(A-F) In situ hybridization stained histological sections for SMC2. SMC2 expression becomes more restricted within epidermal derived feather cells as development progresses. Scale bars 0.1mm.

(G) Diagram of a feather germ showing the position of sections A-F

Figure S3: Changes in AXL and CHL1 expression over time, related to Figure 2

(A-F) In situ hybridization stained histological sections for *CHL1* (A-C) and *AXL* (D-F). Sections on the left are from the basal part of the feather germ and sections to the right are more distal. Scale bars 0.1mm.

(G-H) UMAP feature plots for *CHL1* (G-I) and *AXL* (J-L) from subsetted plucked feather epidermal cells split by samples from H&H stages^8^ 38, 39, and 40 (see Table 1). Red circles highlight increasing numbers of cells for cell types that appear later in feather development (CHL1 – Ramus Medulla, AXL – Ramus Cortex).

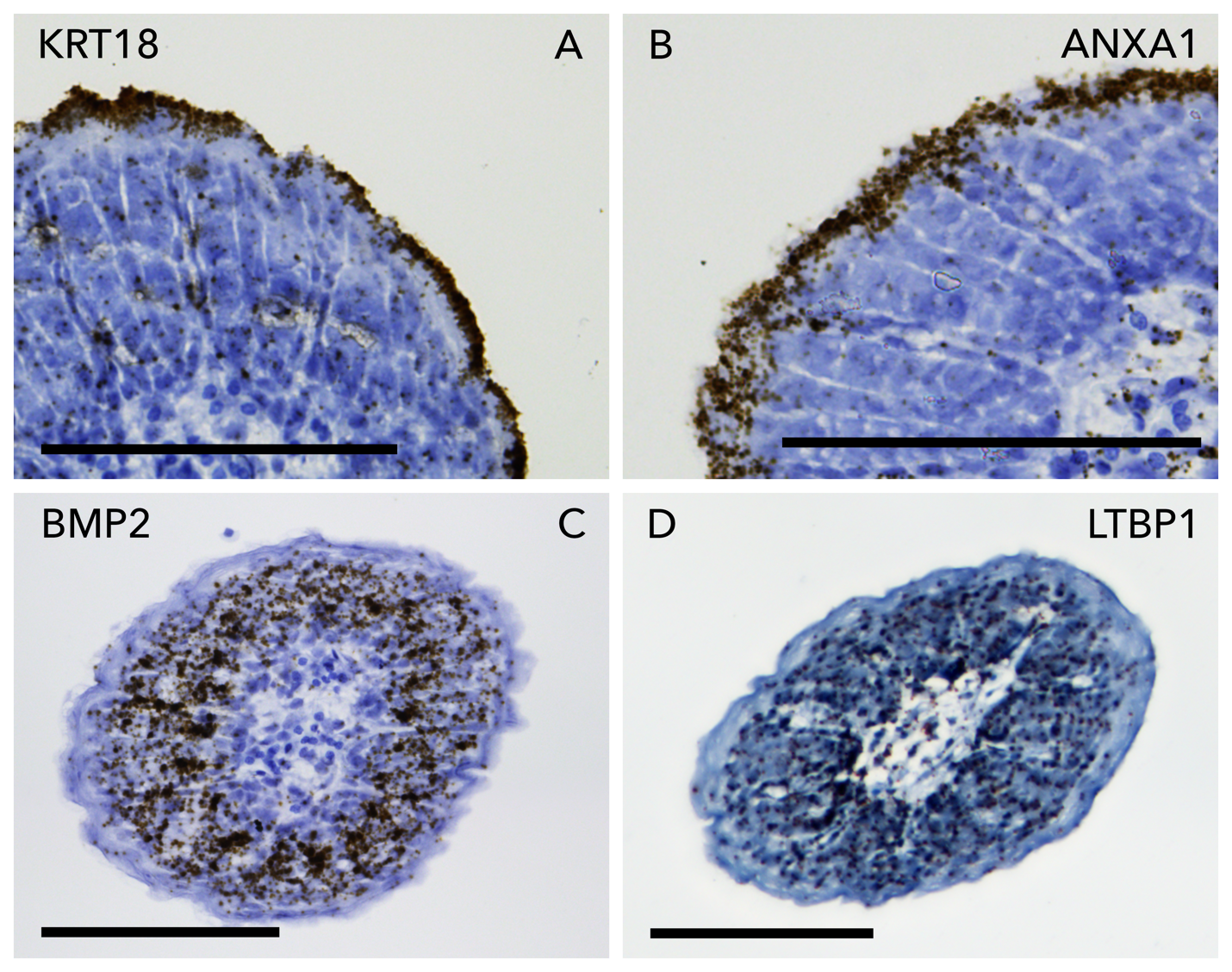

Figure S4: In situ hybridization stained histological sections for genes in need of further elaboration. Scale bars 0.1mm. Related to Figure 2

(A-B) Close ups of KRT18 and ANXA1 staining showing differences in staining within feather sheath. KRT18 stains the periderm sheath to the outside of ANXA1 staining which marks the sheath.

(C) BMP2 staining towards the distal tip of the feather becomes broader and extends beyond the marginal plate.

(D) LTBP1 staining is broad which may indicate an issue with the staining as very restricted expression is predicted by our UMAP (Figure 2BB)
